# Pro-IL-1β activates NF-κB independently of maturation

**DOI:** 10.64898/2026.02.26.708191

**Authors:** Luqiu Mou, Conggang Zhang

## Abstract

Interleukin-1β (IL-1β) is an essential pro-inflammatory cytokine which functions as a key factor in innate immunity. The precursor protein, pro-IL-1β, has long been regarded as an inactive form in innate immune responses. Here, we unveil the biological function and regulation of pro-IL-1β in activating the NF-κB signaling pathway, which is distinct from IL-1β signaling. The expression and release of pro-IL-1β are induced by inflammatory stimuli and then pro-IL-1β acutely activates the gene transcription driven by NF-κB in a dose dependent manner. This activity is resistant to IL-1 receptor antagonist (IL-1Ra). The signal transduction triggered by pro-IL-1β relies on MyD88 and endocytosis. We further demonstrate that the N-terminal pro-peptide primarily contributes to this activity. Furthermore, we identify TLR7 and TLR8 as the binding partners of pro-IL-1β in vitro and as the potential receptors mediating pro-IL-1β-induced NF-κB activation. Collectively, this study sheds light on the unique cytokine function of pro-IL-1β and provides new insights into the functional characterization of pro-cytokines in innate immunity.

## Introduction

Interleukin-1β (IL-1β), which was identified as the first cytokine^1^, plays a central role in regulating innate immune responses. It is originally expressed in the pro-form, named as pro-IL-1β, which is encoded by *IL1B* gene. IL-1β can be produced by various cell types such as immune cells, fibroblasts, or even cancer cells. Among these, studies of IL-1β production are more focusing on innate immune cells, especially myeloid cells. Upon stimulation by stress or cytokines, macrophages and monocytes rapidly initiate NF-κB signaling pathways, for example, TLR4 or TNF-α pathway, to induce the transcription of *IL1B* gene^2–4^. The 31-kDa precursor, pro-IL-1β, is further processed into 17-kDa mature IL-1β by activated Caspase-1during inflammation^5^. Then mature IL-1β is released from the cells by vesicle trafficking or the change of cell membrane permeability^6–12^, which still remains elusive. Secreted IL-1β will activate IL-1R1 and lead to positive feedback on NF-κB activation. As an inflammatory cytokine, IL-1β is identified to be associated with a variety of diseases, such as cancer, stroke, gout and Crohn’ disease^2,13–15^. Up to now, IL-1β is extensively studied and characterized to be involved in many physiological or pathological processes. Also, secreted mature IL-1β is long thought to be the function unit of *IL1B* gene product. However, biological functions of the precursor pro-IL-1β is scarcely studied and understood.

Full-length pro-IL-1β contains 269 aa which is composed of a flexible N-terminal (116 aa) and a C-terminal mature domain (153 aa). The C-terminal mature domain can bind to IL-1R1 and IL-1RAP, forming a heterotrimer and initiate downstream signaling, whereas the N-terminal part is poorly investigated. Compared with mature IL-1β, its pro-form was found to be biologically inactive and needs proteolytic cleavage to convert into the C-terminal mature form^16–18^. Recent studies have been illustrating the potential biological functions of pro-IL-1β. It was reported that deubiquitinase POH1 interacts with pro-IL-1β and inhibits its ubiquitination on K63, then suppresses the maturation of pro-IL-1β^19^. Another research indicated that pro-IL-1β in perfusate is an early-stage biomarker of the donor lung injury during transplantation^20^. An up-to-date study revealed a moonlighting function of pro-IL-1β in head and neck squamous cell carcinoma (HNSCC)^21^. pro-IL-1β was found to be upregulated in HNSCC tissues and cell lines. The mechanistic study suggested that intracellular pro-IL-1β binds to RACK1 and inhibit UBE2T-mediated ubiquitination of RACK1 protein, thereby increases the stability of RACK1 and activates RhoA GTPase signaling in HNSCC cells. This process eventually contributes to cancer invasion and metastasis. Nevertheless, there is still lack of evidence whether pro-IL-1β has a general function under physiological conditions, which is an attractive mystery.

In this study, we discovered pro-IL-1β as a functional cytokine that activates NF-κB signaling pathway. Extracellular pro-IL-1β is more than the precursor of IL-1β and functions independently as an agonist in innate immune responses. Mechanistic studies of pro-IL-1β indicate a distinct mechanism from IL-1β-mediated NF-κB activation. The strong and rapid activation depends on MyD88 and cannot be suppressed by IL-1Ra. Cellular responses of pro-IL-1β stimulation relies on the endocytosis of the protein. Moreover, STING-TBK1 axis is also activated via the overreaction of NF-κB activation. We further focus on the N-terminal pro-peptide of pro-IL-1β and demonstrated it as a functional fragment, even after trypsin digestion. Finally, we identified TLR7/8 as the potential receptors for pro-IL-1β.

## Results

### pro-IL-1β activates NF-κB signaling pathway

The research on functional characterization of pro-IL-1β started from a surprising incident. We accidentally found it as a binder of mouse TLR9 in GST pull-down assay when studied the regulation of TLR pathway (Fig. S1a). We further tested the interaction of human TLR9 and pro-IL-1β in HEK293T cells. As shown in Fig. S1b and Fig. S1c, TLR9-FLAG, UNC93B1 and pro-IL-1β-HA were overexpressed in different combinations. In this Co-IP assay, pro-IL-1β interacted with TLR9 either in the presence or in the absence of UNC93B1, indicating the potential direct interaction. The binding showed no dependence on detergents used in immunoprecipitation. Besides, there was no significant difference between non-reducing and reducing conditions in the assay, which suggested free cysteine residues on pro-peptide of pro-IL-1β didn’t form inter– or intra– molecule disulfide bonds in the complex. The rigorous binding between pro-IL-1β and TLR9 triggered our curiosity and lead to further investigations on the biological function of pro-IL-1β.

We firstly reproduced the release of pro-IL-1β in cell lines and primary cells used in our study (Fig 1a, b, c). Expression of pro-IL-1β was able to be induced by LPS, and the maturation was trigger by nigericin. pro-IL-1β existed both in cytosol and medium after stimulation. This result provided the experimental basis on following treatments with pro-IL-1β, and suggested our treatment as the mimic of physiological condition. We further purified recombinant untagged human IL-1β (Fig. S2a) and pro-IL-1β (Fig. S2b) as the stimuli for these immune cells. Considering toll-like receptors are involve in the activation of both NF-κB and ISG transcription, THP1-Lucia NF-κB cells and THP1-Lucia ISG cells were selected as reporter cell lines for the functional tests. In THP1-Lucia NF-κB cells, pro-IL-1β stimulation showed a significant activation signal, which was stronger than LPS or IL-1β stimulation. And the signal strength didn’t depend on pore-forming cytolysin perfringolysin O (PFO) (Fig. 1d). As in THP1-Lucia ISG cells, pro-IL-1β activated ISRE responses compared with IL-1β, but the signal was weaker than cGAMP treatment (Fig. 1e). Furthermore, we purified recombinant untagged mouse IL-1β (Fig. S3a) and pro-IL-1β mutant (C9S, C34S, C42S, C104S, C116S, Fig. S3b) in order to test the conservation of this activation signal in rodents. N-terminal cysteine-free mutations was introduced into mouse pro-IL-1β to reduce the aggregation of protein during purification. It was shown that pro-IL-1β could activate key regulators in NF-κB signaling pathway in mouse cell lines RAW264.7 (Fig. 1f) and iBMDM (Fig. 1g), which had a similar pattern with LPS stimulation and much stronger than IL-1β. Hence, the prominent activation of NF-κB signaling by pro-IL-1β indicated the potential biological function of this protein.

**Figure 1.**
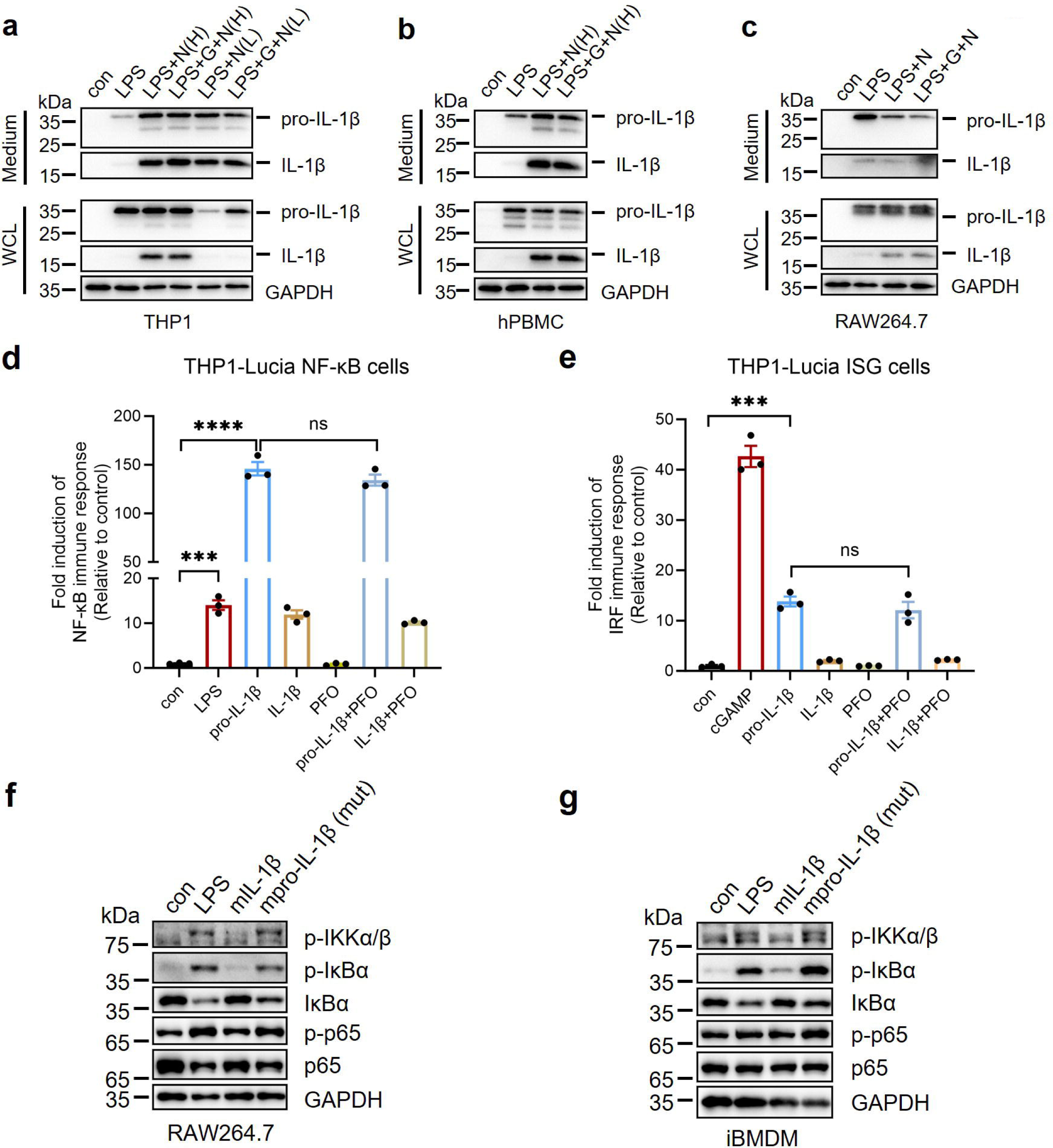
| Characterization of pro-IL-1β as an activator of NF-κB signaling pathway. **a, b, c,** Production and secretion of pro-IL-1β and IL-1β of cell lines used in this study, detected by immunoblotting. THP1 (**a**) and hPBMC (**b**) cells were pre-treated 4 h with 1 μg/mL LPS for pro-IL-1β production, then protected with or without 5 mM glycine (+G) for 1 h. Finally, IL-1β production was stimulated by addition of nigericin (N (H): 5 μM for 30 min; N (L): 0.5 μM for 12 h). For RAW264.7 (**c**), cells were pre-treated 4 h with 1 μg/mL LPS then protected with or without 5 mM glycine (+G) for 1 h. Finally, IL-1β production was stimulated 1 h by addition of 10 μM nigericin. Con: untreated sample; LPS: lipidpolysaccharides; N: nigericin; G: glycine; WCL: whole cell lysate. **d, e,** Activation of THP1-Lucia NF-κB cells (**d**) and THP1-Lucia ISG cells (**e**) with different stimuli. Cells were treated with 1 μg/mL LPS, 1 μM cGAMP, 1 μM pro-IL-1β or 1 μM IL-1β for 16 h. In PFO treated groups, 0.02 μg/mL PFO was added to the medium with or without other stimuli. **f, g,** Activation of NF-κB signaling pathway in mouse-oriented cell lines, RAW264.7 (**f**) and iBMDM (**g**). Cells were treated with 0.5 μM mIL-1β or 0.5 μM mpro-IL-1β (mut) for 1 h, or 1 μg/mL LPS for 1 h. mIL-1β: mouse IL-1β; mpro-IL-1β (mut): mouse pro-IL-1β (C9S, C34S, C42S, C104S, C116S). Error bars represented S.E.M. Unpaired student-t test performed, ns = p>0.05, *** = p ≤0.001, **** = p ≤0.0001.

In order to confirm that pro-IL-1β but not other factors or fragments induced NF-κB activation, we afterwards dissected the origin of the signal. A time-course and concentration gradient of pro-IL-1β stimulation was applied to the reporter assay in THP1-Lucia NF-κB cells (Fig. 2a). The data showed that pro-IL-1β activated NF-κB signaling in a time– and dose-dependent manner. Also, we carefully examined the influence of endotoxin on this assay. The endotoxin level of purified pro-IL-1β was measured at 0.06 EU/μg (Table S1) using standard LAL method. Compared with 1 μg/ml LPS and endotoxin standard, pro-IL-1β showed conspicuous activation signal (Fig. 2b) within a relatively low endotoxin level, and had no synergy with LPS in this assay (Fig. S4a). Additionally, pro-IL-1β didn’t exhibit synergy with R848, another common agonist for NF-κB signaling which activates TLR7/8 (Fig. S4b). The response was rapid and intense although the treatment of pro-IL-1β only lasted 0.5 h (Fig. 2b). Immunoblotting of stimulated cells indicated that pro-IL-1β clearly activated regulators which lead to the activation of NF-κB (Fig. 2c). It was noticed that STING-TBK1 axis was activated as well, which would be discussed in the following experiments. We then tested the stability of pro-IL-1β in our reporter assay system. It was shown by immunoblotting that pro-IL-1β was stable in RPMI 1640 medium or THP1 cells’ medium (Fig. 2d). LC-MS results indicated the intactness of pro-IL-1β and IL-1β in solution (data not shown). Moreover, EC_50_ of NF-κB activation by pro-IL-1β is measured at about 14.33 nM (Fig. 2e, 2f), which highlights the authenticity and intensity of pro-IL-1β stimulation. The studies above showed that the intact pro-IL-1β serves as a unique agonist for NF-κB signaling pathway.

**Figure 2.**
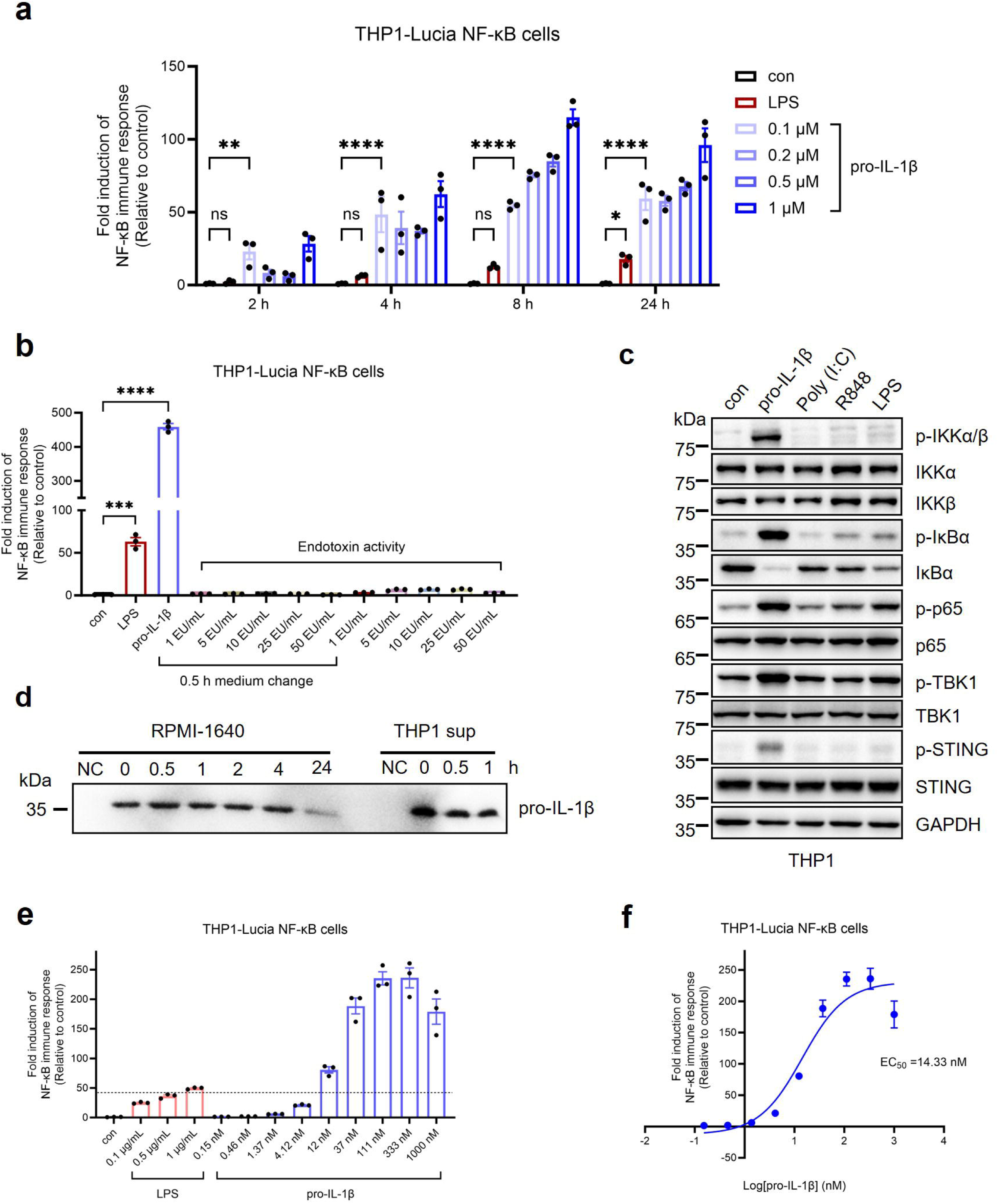
| pro-IL-1β intact protein, but not degraded products, can activates canonical NF-κB signaling pathway in a dose-dependent manner. **a,** Dose– and time-dependence of activation of THP1-Lucia NF-κB cells by pro-IL-1β stimulation. Cells were treated with 1 μg/mL LPS and a concentration gradient of pro-IL-1β. Medium were harvested at different time point for detection. **b,** Comparison of activation level of LPS, pro-IL-1β and endotoxin standard. Stimuli include 0.5 μM pro-IL-1β, 1 μg/mL LPS and a concentration gradient of endotoxin standard (1 EU/mL to 50 EU/mL, 1 ng LPS = 2.5 EU). Medium were harvested after 16 h for detection. For medium change groups, stimuli were removed after 0.5 h treatment. **c,** pro-IL-1β activates canonical NF-κB signaling pathway. THP1 cells were stimulated with 0.5 μM pro-IL-1β, 10 μg/mL poly(I:C), 5 μg/mL R848 or 1 μg/mL LPS for 1 h at 37 □. Cells were lysed and analyzed with immunoblotting. **d,** Medium stability of pro-IL-1β. 0.5 μM pro-IL-1β was incubated with RPMI 1640 medium or THP1 cells’ medium (sup) at 37 □. Medium samples at different time point were analyzed with immunoblotting. NC: untreated groups. **e, f,** Columns (**e**) and non-linear regression curve (**f**) of NF-κB activation by pro-IL-1β. Cells were stimulated with different concentrations of LPS and pro-IL-1β for 16 h. pro-IL-1β was removed after 0.5 h stimulation. Error bars represented S.E.M. Unpaired student-t test performed, ns = p>0.05, * = p ≤0.05, ** = p ≤0.01, *** = p ≤0.001, **** = p ≤0.0001.

### Signal transduction of pro-IL-1**β**-mediated NF-**κ**B activation

The unusual function of pro-IL-1β lead to further exploration on how it activates innate immune responses. Knock-out studies showed that NF-κB activation depended on MyD88 (Fig. 3a), and ISG activation relied on STING (Fig. 3b). Immunoblotting results suggested that the activation of STING-TBK1 axis also had dependence on MyD88, since MyD88^−/−^ impaired the phosphorylation of both proteins mediated by pro-IL-1β stimulation (Fig. 3c). Thus, we speculated that ISG activation results from the over-activation of NF-κB.

**Figure 3.**
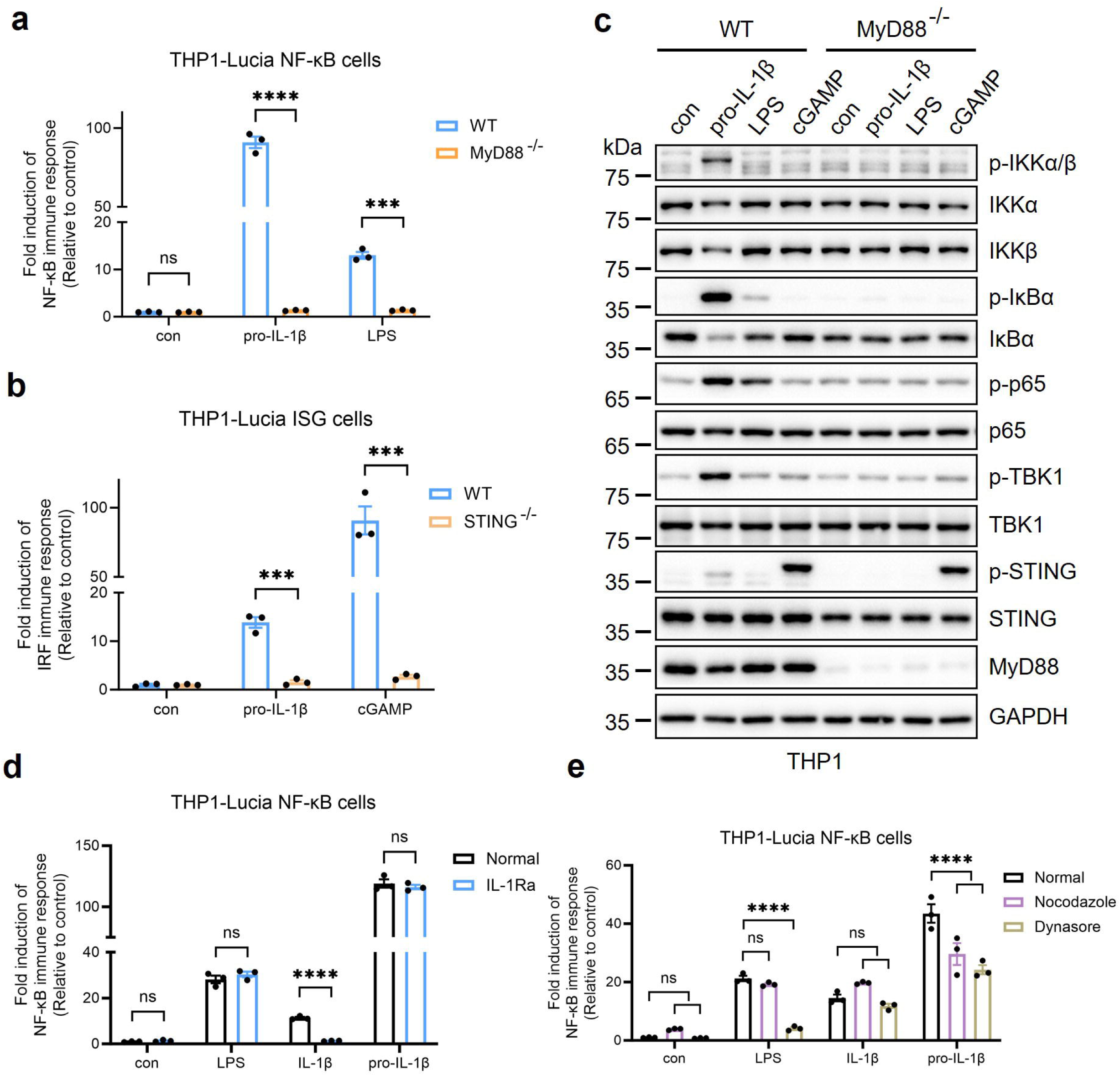
| Biochemical properties of NF-κB activation stimulated by pro-IL-1β. **a, b,** Effects of MyD88^−/−^ (**a**) and STING^−/−^ (**b**) on activation of THP1-Lucia cells by pro-IL-1β. For NF– κB reporter cells, 0.1 μM pro-IL-1β and 1 μg/mL LPS were used for treatment. For ISG reporter cells, 0.5 μM pro-IL-1β and 1 μM cGAMP were used. All of the cells were cultured for 16 h at 37 □. pro-IL-1β was removed after 0.5 h stimulation. Error bars represented S.E.M. Two-way ANOVA (**a**) and unpaired student-t test (**b**) performed, ns = p>0.05, *** = p ≤0.001, **** = p ≤0.0001. **c,** Immunoblotting analysis of activation of NF-κB and cGAS-STING pathways by different stimuli in WT and MyD88^−/−^ THP1 cells. Cells were treated with 0.5 μM pro-IL-1β, 1 μg/mL LPS, or 1 μM cGAMP for 1 h at 37 □, then lysed for analysis. WT: wild-type. **d,** Effects of IL-1Ra on NF-κB activation. Cells were pre-treated with 1 μM IL-1Ra for 30 min, then stimulated by 1 μg/mL LPS, 0.1 μM IL-1β or 0.1 μM pro-IL-1β. Medium was harvested after 16 h for analysis. Cytokines were removed after 0.5 h stimulation. Two-way ANOVA (con, LPS and pro-IL-1β) and unpaired student-t test (IL-1β) performed, ns = p>0.05, **** = p ≤0.0001. **e,** Effects of endocytosis inhibitors on NF-κB activation. Cells were pre-treated with 20 μM nocodazole or 10 μM dynasore for 30 min, then stimulated by 1 μg/mL LPS, 0.1 μM IL-1β and 0.1 μM pro-IL-1β for 16 h. Two-way ANOVA performed. ns = p>0.05, **** = p ≤0.0001.

Given that IL-1β activates downstream signaling through MyD88, we subsequently tested whether pro-IL-1β shares the same pathway with IL-1β during signal transduction. It was concluded that the inhibitory protein IL-Ra could not abolish pro-IL-1β-mediated NF-κB activation, whereas IL-1β signaling was almost totally inhibited (Fig. 3d). And the study on endocytosis inhibitors showed that NF-κB activation by pro-IL-1β could be inhibited by nocodazole and dynasore, which was different from IL-1β as well (Fig. 3e). Since previous studies reported pro-IL-1β doesn’t interact with IL-1R1^22,23^, it exerted a distinct transduction and regulation properties.

Both pro-IL-1β and IL-1β mediated NF-κB activation, but pro-IL-1β functioned independently without any correspondence to IL-1β production. The differences in signal intensity, regulation and mechanism of action emerged to be important. We further analyzed the downstream gene regulation of NF-κB activation driven by pro-IL-1β and IL-1β using RNA-seq. The venn diagram exhibited that 2293 genes were modulated by pro-IL-1β stimulation compared with negative control, while the modulation induced by IL-1β stimulation is much weaker (Fig. S5a). Together with volcano plot (Fig. S5b), pro-IL-1β significantly regulated 488 genes in contrast of IL-1β, where 380 genes were up-regulated and 108 genes were down-regulated, as presented in Table S2. And it was found that *IL1B* was involved in the up-regulation cluster, which suggested the existence of positive feedback in signal transduction of the pathway. Moreover, GO term analysis (Fig. S5c) showed that pro-IL-1β stimulation might re-shape the expression pattern of secreted proteins and membrane receptors. And the downstream signals of pro-IL-1β was probably related to signal cascade of immune responses.

### Pro-peptide of pro-IL-1**β** plays a major role in NF-**κ**B activation

Since pro-IL-1β exhibited distinct function from IL-1β, it was naturally suggested that the pro-peptide might be essential for the activity of pro-IL-1β. We then tried to purify the N-terminal (1-116) of pro-IL-1β for further functional characterization. As shown in Fig. 4a, a drICE cutting site (DEVD/A) was introduced between D116 and A117 in order to avoiding severe degradation and obtaining intact pro-peptide. We employed the dual-proteolytic strategy using GST-HRV3C-pro-peptide-drICE-IL-1β construct, which led to successful acquisition of pro-peptide via 3C protease and drICE co-digestion (Fig. 4b). It could be purified using ion-exchange chromatography and gel-filtration, and the intactness was identified by LC-MS.

**Figure 4.**
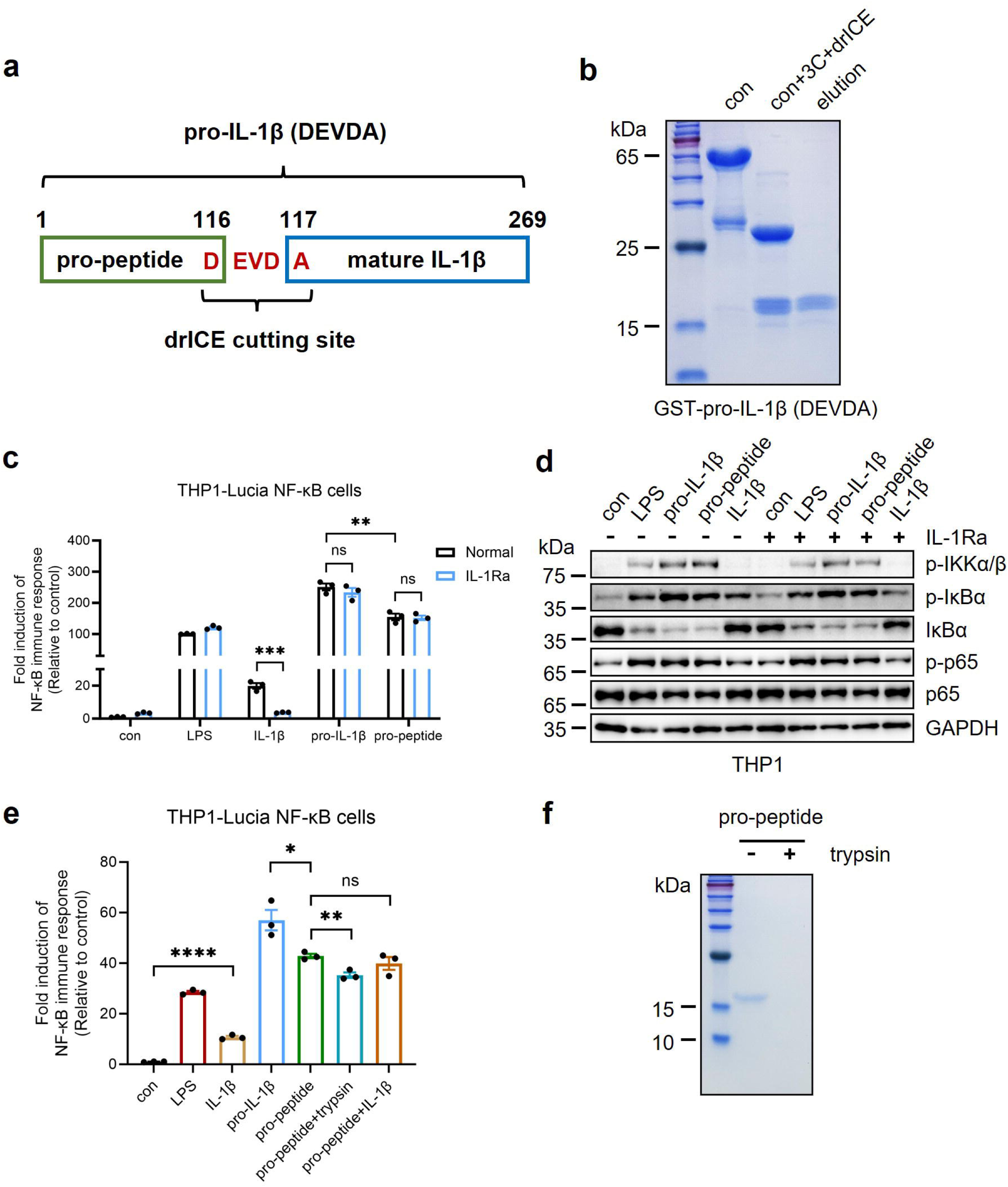
| pro-peptide of pro-IL-1β activates NF-κB signaling pathway. **a,** Construct design of drICE-cleavable pro-IL-1β. **b,** Biochemical characterization of cleaved pro-IL-1β. **c, d,** pro-peptide activates NF-κB signaling pathway. Reporter assay (**c**) and immunoblotting analysis (**d**) of THP1 cells after stimulation. Cells were stimulated by 1 μg/mL LPS, 0.1 μM IL-1β, 0.1 μM pro-IL-1β or 0.1 μM pro-peptide for 16 h (**c**). In IL-1Ra groups, cells were pre-inhibited by addition of 1 μM IL-1Ra. For immunoblotting analysis, treated cells were harvested and lysed immediately after 1 h stimulation with 1 μg/mL LPS, 0.5 μM IL-1β, 0.5 μM pro-IL-1β or 0.5 μM pro-peptide (**d**). **e,** Characterization of functional fragments in pro-peptide. THP1-Lucia NF-κB cells were treated with 0.1 μM IL-1β, 0.1 μM pro-IL-1β or 0.1 μM pro-peptide for 0.5 h, or 1 μg/mL LPS for 16 h. All treatments were detected after 16 h incubation. pro-peptide+trypsin: 0.1 μM pro-peptide was pre-treated with 1:10 (w/w) trypsin at 4 for 3 h. pro-peptide+IL-1β: 0.1 μM pro-peptide was pre-incubated with 0.1 μM IL-1β at 4 for 1 h. **f,** SDS-PAGE analysis of undigested and digested pro-peptide. Error bars represented S.E.M. Unpaired student-t test performed, ns = p>0.05, * = p ≤0.05, ** = p ≤0.01, *** = p ≤0.001, **** = p ≤0.0001.

The function of purified pro-peptide was examined by reporter assay (Fig. 4c) and immunoblotting (Fig. 4d). Consistent with our speculation, pro-peptide activated NF-κB signaling and couldn’t be suppressed by IL-1Ra. Both signals induced by pro-peptide and pro-IL-1β was stronger than LPS-induced one. To our surprise, the pro-peptide didn’t maintain full activity compared with full-length protein, and couldn’t be rescued by co-stimulation together with mature IL-1β (Fig. 4e). In addition, the activity of pro-peptide was trypsin-resistant (Fig. 4e, f) whereas there was still a minor decrease of the activity after trypsin digestion. The intriguing phenomenon indicated a complicated molecular mechanism of NF-κB activation by pro-IL-1β. Nevertheless, we anchored functional motif(s) in trypsin-treated N-terminal fragments of pro-IL-1β, which provided hints for further studies.

### TLR7/8 are potential receptors for pro-IL-1**β**

The validation of function of pro-IL-1β brought us back to the original finding where pro-IL-1β interacted with mouse TLR9 in vitro. Our results indicated that TLRs was supposed to be the receptors for pro-IL-1β. Sequence alignment of human TLRs suggested that TLR3/7/8/9, instead of other TLRs, were noteworthy targets for further investigation (Fig. S6a). We then purified recombinant full-length TLR3/7/8/9 and Z-loop cleaved TLR7/8/9 (Fig. S6b). The proteins were applied to GST-pull down assay (Fig. 5a). In comparison to GST and GST-tagged IL-1β, GST-pro-IL-1β rigorously bound to purified TLRs in vitro. Also, we examined whether the interactions depended on the regulatory protein UNC93B1 in contrast with another important cytokine precursor in inflammation, pro-IL-18 (Fig. S7a). GST-pro-IL-1β was able to bind TLR3/7/8/9 regardless of UNC93B1, while GST-pro-IL-18 rarely bound to TLRs. A more detailed Co-IP study also showed the interactions of pro-IL-1β and TLR3/7/8/9 in cell-based system (Fig. S7b). The studies above provided evidences that TLR3/7/8/9 were binders of pro-IL-1β and could be potential receptors for pro-IL-1β stimulation.

**Figure 5.**
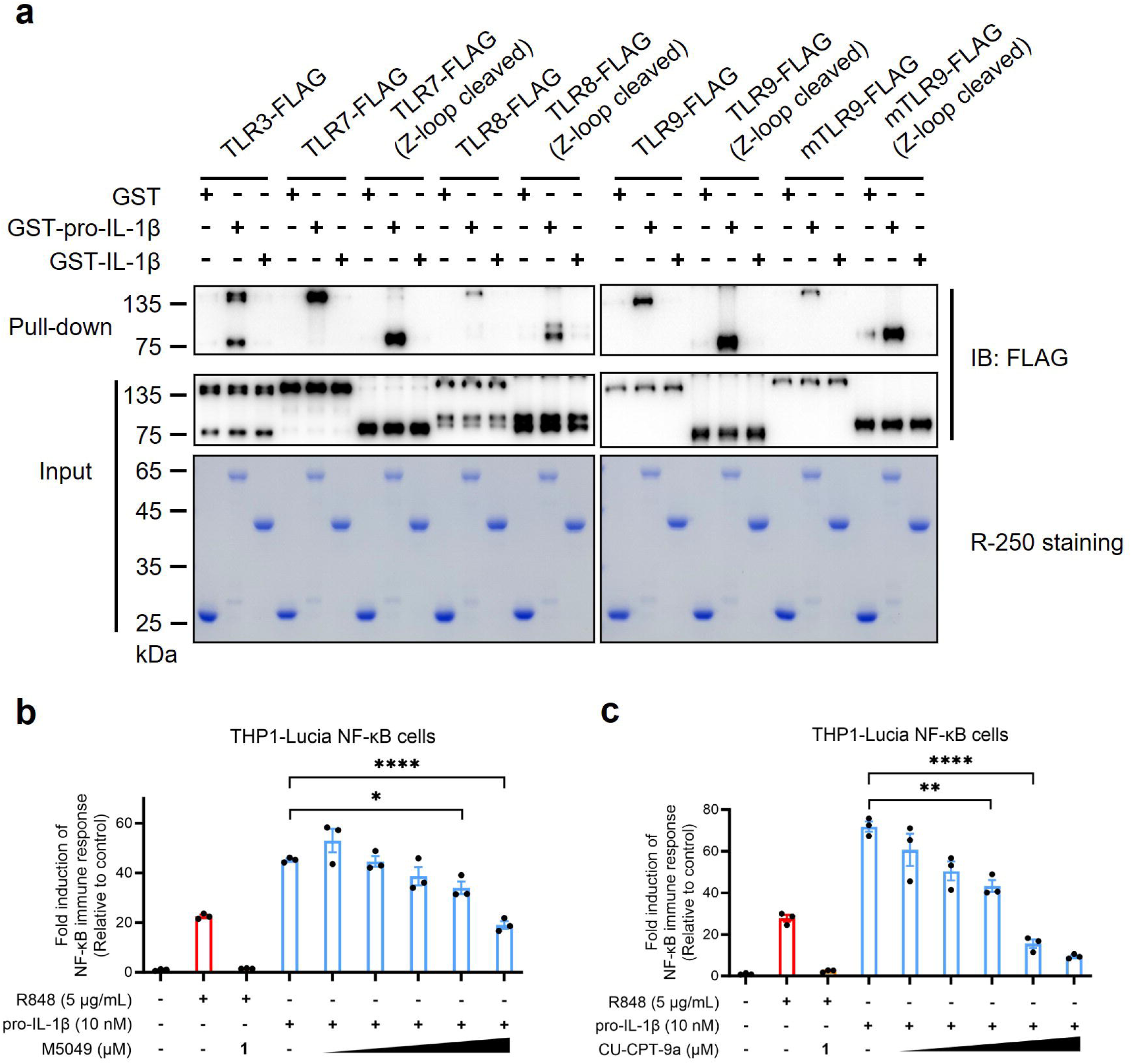
| TLR7/8 serve as potential receptors for pro-IL-1β. **a,** pro-IL-1β binds to TLR3/7/8/9 in vitro. Purified TLRs protein was incubated with GST-tagged cytokines for GST pull-down assay. TLRs were co-expressed with UNC93B1 for better yields. A drICE cutting site (DEVDA) was introduced to Z-loop of TLR to obtain a drICE-cleavable construct, in order to mimic Z-loop cleaved TLR by drICE cleavage. Samples were analyzed with immunoblotting and Coomassie staining. mTLR9: mouse TLR9; controls: GST and GST-IL-1β. **b, c,** Effects of TLR7/8 inhibitors on NF-κB activation. Cells were pre-inhibited by 2-50 μM M5049 (**b**) or CU-CPT-9a (**c**) for 30 min, then activated by 10 nM pro-IL-1β. Positive control groups were treated with 5 μg/mL R848. Error bars represented S.E.M. Unpaired student-t test performed, ns = p>0.05, * = p ≤0.05, ** = p ≤0.01, **** = p ≤0.0001.

Since previous experiment indicated that MyD88^−/−^ abolished the activation of NF-κB by pro-IL-1β, we speculated that TLR3 might not be involved in the signal transduction. Likewise, TLR9 was excluded due to the failure of ODN2006 activation in THP1-Lucia NF-κB cells according to the manufacturer’s instructions. Thus, we focused on the study of TLR7 and TLR8, which were identified as ssRNA receptors in innate immune responses. Inhibitory studies found that M5049^24^ (Fig. 5a) and CU-CPT-9a^25^ (Fig. 5b) suppressed pro-IL-1β-mediated NF-κB activation in a dose-depended manner. However, these inhibitors showed limited effects. 1 μM inhibitors were sufficient for the inhibition of 15.9 μM R848, while 10 μM concentration was not enough to eliminate the signal of 10 nM pro-IL-1β, suggesting a non-competitive inhibition mechanism. The results suggested that TLR7/8 were potential receptors for pro-IL-1β but the binding site could be different from canonical agonists or antagonists.

## Discussion

As a brief summary, we unveiled pro-IL-1β as a functional cytokine that activates NF-κB signaling pathway. In the proposed working model (Fig. 6), the production of pro-IL-1β is driven by upstream stresses like LPS and IL-1β. It will be directly released from cells, or processed by active Caspase-1 during inflammation. Previous study had shown that pro-peptide is also released to extracellular side^26^. pro-IL-1β then activates downstream signaling which depends on MyD88 and cannot be inhibited by IL-1Ra. The signal transduction is associated with endocytosis and suppressed by TLR7/8 inhibitors, suggesting the potential of TLR7/8 as the receptors. Transcription of genes including *IL1B* are rapidly and strongly induced by NF-κB activation, and it is proposed to serve as positive feedback of the signaling cascade.

**Figure 6.**
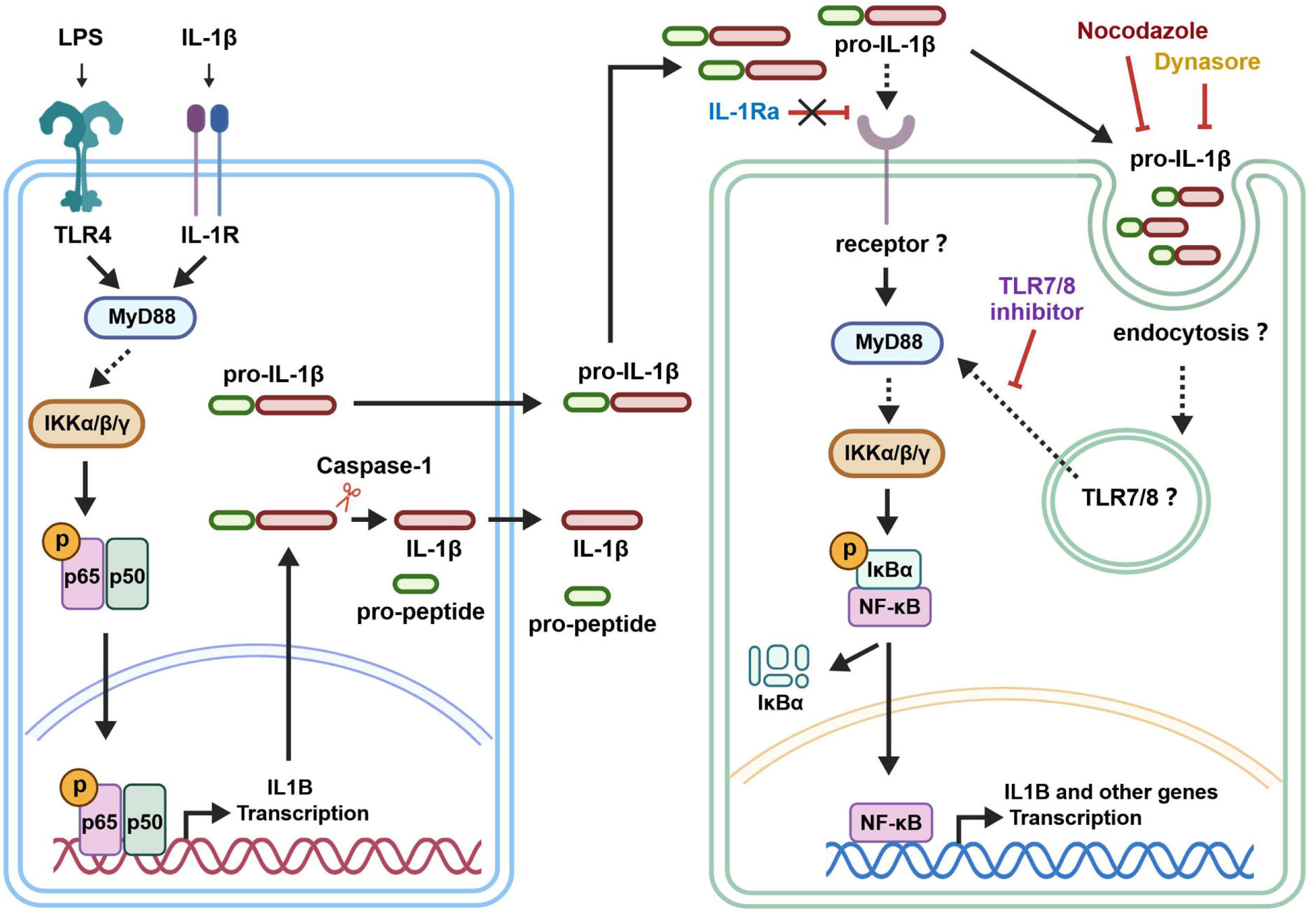
| Updated working model of NF-κB activation by pro-IL-1β. LPS or IL-1β activates NF-κB signaling pathway. Upon activation, *IL1B* transcription is induced and leads to the high-level expression of pro-IL-1β protein. Inflammasome-activating Caspase-1 cleaves pro-IL-1β and mature IL-1β is released from the cell. In physiological conditions, pro-IL-1β is also a functional cytokine that can be independently released from cells and activates NF-κB signaling pathway, which has distinct features from IL-1β. The working model was created with BioGDP.com.

The current study found that pro-IL-1β rigorously activated NF-κB signaling pathway in myeloid cells. And the response to pro-IL-1β stimulation was rapid, intense and sensitive compared with LPS and IL-1β. The EC_50_ value of pro-IL-1β reached around 14.33 nM. We infer that pro-IL-1β plays a predominant role in signal amplification during inflammation. Monocytes and macrophages receiving the stress would express and release pro-IL-1β to extracellular region, and pro-IL-1β functions across the cells as a ‘messenger’ so that the stimuli could be promptly responded. Cell death was observed when treated with high concentration of pro-IL-1β, as well for long-time treatment, which is consistent with the indication. Our data unprecedentedly uncovered pro-IL-1β is a functional cytokine rather than a precursor protein in innate immune responses.

We further analyzed how NF-κB signaling induced by pro-IL-1β is regulated. Distinct from IL-1β, pro-IL-1β activated downstream signaling which couldn’t be suppressed by IL-1Ra, and the activation relied on endocytosis, although it showed the MyD88 dependence. The results suggested that there should be a unique mechanism for activation and regulation of pro-IL-1β. Failure of IL-1Ra inhibition moved our attention to the N-terminal pro-peptide. The current study showed that pro-peptide of pro-IL-1β reserved most of the activity and functioned independently. Further trypsin challenge didn’t lead to the complete loss of activity. On the contrary, over half of the activity remained in comparison of full-length protein. We speculated that the minimal functional motif of pro-IL-1β resides in trypsin-resistant peptides of pro-peptide and full activity requires intactness of the protein. However, the partial loss of activity observed in this assay indicated there might be inter– or intra-domain interactions of pro-peptide, which is sophisticated and needs further investigation.

In another aspect, correlations between TLRs and pro-IL-1β can not be ignored as we started this research from the observation of pro-IL-1β binding to mouse-TLR9. We initially selected TLR3/7/8/9 on the basis of sequence identity. Also, the rich of acidic residues on pro-peptide of pro-IL-1β might be potential mimick of oligonucleotides ligands for these TLRs, which supported our rationale. GST-pull down and Co-IP assays showed that pro-IL-1β interacted with TLR3/7/8/9. Based on the feature of signaling and cell line we used in the study, as mentioned in the results, M5049 and CU-CPT-9a were applied to the inhibitory experiments. It was found that the two inhibitors suppressed pro-IL-1β mediated NF-κB activation. But to our surprise, the inhibition was not strong enough compared with the one of R848. The result led to another speculation that if pro-IL-1β bound to TLRs in an uncharacterized binding site different from natural ligands and inhibitors, therefore the inhibitors couldn’t work efficiently. In addition, we tried to knock out TLR7 or TLR8 in our reporter assay. But unfortunately, it was too difficult to obtain successful KO THP1 cells even after six sgRNAs for each gene were used. So we only defined TLR7/8 as the potential receptors for pro-IL-1β which needs more structural and biochemical studies.

Last but not least, the potential of pro-IL-1β as a therapeutic target is an intriguing future direction. Inhibition of IL-1β signaling by recombinant inhibitory protein, ligand trap or antibody was proved to be potent in different pathological conditions^27,28^. Anakinra, a engineered human IL-1Ra, showed potency in rheumatoid arthritis (RA), cryopyrin-associated periodic syndrome (CAPS) and neonatal-onset multisystem inflammatory disorder (NOMID)^29^. And the monoclonal antibodies like Canakinumab, which was extensively studied in clinical trials, was approved for treatment in RA, ankylosing spondylitis (AS) and gout. Considering the notable function of pro-IL-1β in contrast with IL-1β, we suggest pro-IL-1β to be a noteworthy target for drug development in inflammatory diseases. In vivo studies are demanded for conditions that pro-IL-1β is involved in.

In conclusion, we demonstrated that pro-IL-1β serves as a functional cytokine in innate immunity and strongly activates NF-κB signaling pathway. Investigations on signal modulation showed that the activation depends on MyD88 and endocytosis, and can’t be inhibited by IL-1Ra. The function is mainly contributed by N-terminal pro-peptide. Biochemical studies identify TLR7/8 as the potential receptors for pro-IL-1β. Our study illustrates the essential biological function of pro-IL-1β, which was thought to be inactive in innate immune responses.

## Acknowledgements

The study is also supported by the National Natural Science Foundation of China (project approval number: 32571031 to C.G. Zhang), and grants from Tsinghua-Peking Center for Life Sciences, Chongqing Key Laboratory of Precision Diagnosis and Treatment for Kidney Diseases, Beijing Frontier Research Center for Biological Structure, and Tsinghua University Initiative Scientific Research Program and Dushi program.

## Author contributions

C.Z. conceived the project. L.M. and C.Z. designed all experiments. L.M. performed the experiments. L.M. and C.Z. analyzed the data and wrote and manuscript.

## Author Information

The authors declare no competing financial interests. Correspondence and requests for materials should be addressed to C.Z.

## Materials and methods

KEY RESOURCES TABLE.

KEY RESOURCES TABLE
| REAGENT or RESOURCE | SOURCE | IDENTIFIER |
| --- | --- | --- |
| **Antibodies** |  |  |
| Anti-phospho-IKKα/$\beta$ (Ser176/180) (16A6) | Cell Signaling Technology (CST) | 2697 |
| Anti-IKKα (D3W6N) | CST | 61294 |
| Anti-IKKβ (D30C6) | CST | 8943 |
| Anti-phospho-IκBα (Ser32/36) (5A5) | CST | 9246 |
| Anti-IκBα | CST | 9242 |
| Anti-phospho-NF-κB p65 (Ser536) (93H1) | CST | 3033 |
| Anti-NF-κB p65 (D14E12) | CST | 8242 |
| Anti-phospho-STING (Ser366) (D7C3S) | CST | 19781 |
| Anti-STING (D2P2F) | CST | 13647 |
| Anti-phospho-TBK1 (Ser172) (D52C2) | CST | 5483 |
| Anti-TBK1 | Abcam | ab40676 |
| Anti-MyD88 (D80F5) | CST | 4283 |
| Anti-GAPDH | Servicebio | GB15004 |
| Anti-β-Tubulin | Abmart | M20005 |
| Anti-UNC93B1 | Proteintech | 28359-1-AP |
| Anti-hpro-IL-1β (human p31) | R&D | MAB201 |
| Anti-hIL-1β (human p17) | CST | 83186 |
| Anti-mpro-IL-1β (mouse p31) | R&D | BAF401 |
| Anti-mIL-1β (mouse p17) | CST | 63124 |
| Anti-DYKDDDDK tag | Sigma | F1804 |
| Anti-HA tag | Abmart | M20003 |
| Rabbit Anti-Goat IgG H&L, HRP conjugated | Bioss | bs-0294R-HRP |
| Goat Anti-Mouse IgG H&L, HRP conjugated | Bioss | bs-0296G-HRP |
| Goat Anti-Rabbit IgG H&L, HRP conjugated | Bioss | bs-0295G-HRP |
| **Cell lines** |  |  |
| HEK293T | ATCC | CRL-3216 |
| Expi293F | Gibco | R79007 |
| THP1 | ATCC | TIB-202 |
| THP1-Lucia NF-κB Cells | Invivogen | thpl-nfkbv2 |
| THP1-Lucia ISG Cells | Invivogen | thpl-isg |
| Human PBMC | YAYUBIO | CZPB025M |
| RAW264.7 | ATCC | SC-6003 |
| iBMDM | This paper | N/A |
| MyD88<sup>-/-</sup> THP1-Lucia NF-κB Cells | This paper | N/A |
| STING<sup>-/-</sup> THP1-Lucia ISG Cells | This paper | N/A |
| **Bacterial strains** |  |  |
| *E. coli* DH5α | This paper | N/A |
| *E. coli* BL21-RIL (DE3) | This paper | N/A |
| **Reagents** |  |  |
| Polyethylenimine (PEI) | Polysciences | 23966-2 |
| Fetal Bovine Serum (FBS) | Excell | FSP500 |
| Dulbecco's modified Eagle's medium (DMEM) | Gibco | 11995500BT |
| RPMI 1640 medium | Gibco | 11875500BT |
| Penicillin-Streptomycin | Solarbio | P1400 |
| Opti-MEM | Gibco | 22600050 |
| Puromycin | Beyotime | ST551 |
| Blasticidin S | Beyotime | ST018 |
| PFO (Perfringolysin O) | This paper | N/A |
| 2'3'-cGAMP | This paper | N/A |
| LPS | Sigma | L2630 |
| Poly(I:C) | Invivogen | tlrl-picw |
| R848 | MCE | HY-13740 |
| NP-40 | Beyotime | ST2045 |
| DDM | Macklin | D806359 |
| GDN | Anatrace | GDN101 |
| DTT | Aladdin | D104860 |
| PMSF | Sangon Biotech | A100754 |
| Aprotinin | Sangon Biotech | A600153 |
| Pepstatin A | Sangon Biotech | A610583 |
| Leupeptin | Sangon Biotech | A600580 |
| β-ME | Solarbio | M8210 |
| Glutathione Sepharose 4B | Cytiva | 17075605 |
| Anti-FLAG M2 Affinity gel | Sigma | A2220 |
| HA-Nanoab-Agarose Beads | Lablead | HNA-25-500 |
| StrataClean resin | Agilent | 400714 |

| Endotoxin detection kit | BIOENDO | KC64T |
| --- | --- | --- |
| **Others** |  |  |
| 0.45 μm PVDF | Millipore | IPVH00010 |
| SOURCE 15Q | Cytiva | 17094705 |
| Superdex 200 Increase 10/300 GL | Cytiva | 28990944 |
| Superose 6 Increase 10/300 GL | Cytiva | 29091596 |

## Cell culture and transient transfection

HEK293T, RAW264.7 and iBMDM cells were cultured in DMEM containing 10% (v/v) FBS and antibiotics (100 U/ml penicillin, 0.1 mg/ml streptomycin). THP1 cells and human PBMCs were cultured in RPMI 1640 medium supplemented with 10% (v/v) FBS, antibiotics (100 U/ml penicillin, 0.1 mg/ml streptomycin) and 0.05 mM β-mercaptoethanol (β-ME). These cell lines for in vitro assays were cultured at the condition of 37 □, 95% humidity and 5% CO_2_.

For transient expression in HEK293T, the transfection mix was prepared using 6 μg plasmid and 18 μg PEI in 1 mL opti-MEM. Plasmids were added equivalently in mass in the condition of co-transfection. After 10 min incubation at RT, the mix was then added to a 10-cm dish with ∼80% cell coverage. Cells were harvested for further experiments after 48 h transfection.

## Constructs and Cloning

Human pro-IL-1β (1-269, Uniprot: P05184), human IL-1β (117-269), mouse pro-IL-1β (1-269, Uniprot: P10749), mouse IL-1β (118-269) and human pro-IL-18 (1-193, Uniprot: Q14116) were subcloned into pGEX-6p-1 vector for recombinant expression. In order to purify pro-peptide of human pro-IL-1β, a drICE cutting site (DEVDA) was introduced between D116 and A117 using the strategy of PCR-based site-directed mutagenesis. To obtain well-behaved mouse pro-IL-1β protein, mutations of N-terminal cysteines (C9S, C34S, C42S, C104S, C116S) were introduced into DNA sequence via PCR. For transient expression in HEK293T, full-length human pro-IL-1β, pro-IL-18 and pro-IL-33 (Uniprot: O95760) were cloned into pCAG vector with a C-terminal hemagglutinin (HA) tag. Full-length human TLR3 (Uniprot: O15455), TLR7 (Uniprot: Q9NYK1), TLR8 (Uniprot: Q9NR97), TLR9 (Uniprot: Q9NR96) and mouse TLR9 (Uniprot: Q9EQU3) were clone into a C-terminal FLAG-tagged pCAG vector. To generate Z-loop cleavable constructs, drICE cutting site was inserted after S440 of hTLR7, S440 of hTLR8, A440 of hTLR9 and E440 of mTLR9. Full-length human UNC93B1 (Uniprot: Q9H1C4) followed with a C-terminal 8xHis tag was cloned into pCAG vector. All constructs above were generated using PCR and homologous recombination.

Annealing was use for generation of clones for CRISPR knockout. Briefly, annealed sgRNAs were inserted into linearized pLentiCRISPR v2 vector using T4 ligase. sgRNAs: MyD88 (5’-CTCCCCTAGGTGCCGCCGGA-3’), STING (5’– ACTCTTCTGCCGGACACTTG-3’).

## Protein Expression and Purification

Human and mouse pro-IL-1β, IL-1β and human pro-IL-18 were recombinantly expressed in BL21 (DE3) *E.coli* strain. Transformed *E. coli* was cultured in LB medium containing 0.1 mg/ml ampicillin at 37 □, 220 rpm. Cultures grown to OD_600_ of 1.0 were induced by 0.2 mM Isopropyl β-D-thiogalactoside (IPTG). After overnight induction at 18 □, cells were harvested by centrifugation at 3,000 g and resuspended in lysis buffer containing 25 mM Tris pH 8.0 and 150 mM NaCl. The resuspension was then supplemented with 1 mM PMSF and applied to sonication. Cell debris was removed by 20,000 g centrifugation for 1 h at 4 □. And the supernatant was incubated with GS4B resin for another 1 h at 4 □. The resin was rinsed four times with lysis buffer. For GST-tagged protein purification, the protein was eluted with lysis buffer plus 25 mM Tris 8.0 and 10 mM GSH. As for untagged protein, 1:100 (w/w) GST-HRV 3C protease was added to resuspended resin. To obtain pro-peptide, 1:200 (w/w) in-house prepared drICE was supplemented together with 3C protease. After overnight digestion at 4 □, protein was eluted with lysis buffer by gravity flow. The eluate was 5-fold diluted with Buffer A (25 mM Tris pH 8.0 and 2 mM DTT) and further purified using SOURCE 15Q. Protein was then eluted from anion exchange chromatography with 20 CV linear gradient of 0-1 M NaCl. Outlet flowthrough (IL-1β only) or fractions containing target protein were concentrated using Amicon Ultra-15 centrifugal filters and applied to Superdex 200 Increases pre-equilibrated with HEPES buffer containing 20 mM HEPES pH 7.5 and 150 mM NaCl. Peak fractions from gel-filtration were analyzed with SDS-PAGE, pooled and flash frozen in liquid nitrogen for further use.

For TLR proteins, plasmids were co-transfected with UNC93B1 in Expi293F cells. In brief, 0.5 mg TLR plasmid and 0.5 mg UNC93B1 plasmid combined with 3 mg PEI were mixed in Opti-MEM and incubated 15 min at RT. The transfection mix was freshly prepared for 1 L cell culture and applied when cell density reached 1.5∼2×10^6^/mL. After 72 h transfection, cells were centrifuged at 1,500 g for 20 min at 4 and resuspended with lysis buffer supplemented with 0.8 μM aprotinin, 2 μM pepstatin A and 5 μg/mL leupeptin. TLR-UNC93B1 complex was then extracted from the cells by 1.5 h incubation with 1% (w/v) N-dodecyl-β-D-maltoside (DDM) at 4 □. Cell debris was pelleted by 30 min centrifugation at 50,000 g and 4 □. The supernatant was incubated with Anti-FLAG M2 Affinity gel for 2 h at 4 □. The resin was washed with wash buffer which consists of 20 mM HEPES pH 7.5, 200 mM NaCl and 0.02% DDM, and the protein complex was eluted with wash buffer plus 1 mg/mL DYKDDDDK peptide. The eluate was concentrated and applied to Superose 6 Increase pre-equilibrated with MES buffer (25 mM MES pH 5.8, 200 mM NaCl and 0.02% DDM). To generate Z-loop cleaved TLRs, 1:200 (w/w) drICE was used for overnight digestion at 4 before gel-filtration. Peak fractions were subsequently analyzed and collected as above.

## Endotoxin measurement

Endotoxin levels of purified pro-IL-1β, IL-1β and pro-peptide were measured with standard LAL method. The endotoxin standards were prepared and diluted with endotoxin-free water into a gradient of 50, 5, 0.5, 0.05 and 0.005 EU/mL for calibration. The standards and samples were mixed with freshly dissolved reaction mix in an equal volume, and the absorbance at 405 nm was measured at 37 by plate reader. Onset OD in data processing was set to 0.2. Endotoxin levels of the samples were calculated based on the standard curve which was linearly regressed with the equation of lgt = algC + b, t for onset time and C for endotoxin level. The measurement unit was subsequently converted to EU/μg for better presentation.

## Detection of pro-IL-1**β** and IL-1**β** release

THP1 (1.5×10^6^ cells), hPBMC (4×10^6^ cells) and RAW264.7 (4×10^6^ cells) cells were seeded per 24-well. The treatment conditions were described as in Figure 1. After stimulation, 1% (v/v) StrataClean resin was added to the medium for protein enrichment at 4 overnight, with protease inhibitor (0.8 μM aprotinin, 2 μM pepstatin A and 5 μg/mL leupeptin) supplemented. Cells were lysed with RIPA buffer (20 mM Tris pH 7.4, 150 mM NaCl, 10% (v/v) glycerol, 1% (w/v) Triton X-100, 0.1% (w/v) SDS, 1 mM EDTA, 1 mM Na_3_VO_4_ and 25 mM sodium β-glycerophosphate). The cell debris was removed by 20,000 g centrifugation at 4 □, and the supernatant was mixed with 5x SDS-loading buffer for analysis.

## Reporter Assay

THP1-Lucia cells were aliquoted as 1×10^5^ per 96-well. Then stimuli were added to the culture and the detailed treatment conditions were described as in figure legends. After stimulation and incubation at 37 □, 20 μL medium from sample wells was rapidly and thoroughly mixed with 50 μL fresh-diluted 0.5 μM coelenterazine, then the luciferase signal was detected by plate reader. Data of experimental groups was normalized to negative control. Non-linear regression was used for EC_50_ measurement of pro-IL-1β stimulation. The equation for curve fitting is Y=Bottom + (Top-Bottom)/(1+10^(LogEC50-X)), X for Log [Agonist] and Y for response. Top and Bottom are plateaus in the units of the Y axis.

## Immunoblotting

Protein samples, beads or cells were mixed, extracted or lysed with 2x or 5x SDS-loading buffer, then resolved on 10% (for proteins over 100 kDa)/15% (for proteins below 100 kDa) polyacrylamide gels. Then proteins were transferred to 0.45 μm PVDF membrane by wet-blotting. The membranes were blocked by 7% fat-free milk in TBST for 1 h at RT, incubated with primary antibodies overnight at 4 and finally incubated with HRP-conjugated secondary antibodies for 1 h at RT. Three 5-min cycles of wash were inserted between the steps above. After another three 10-min washes to remove the secondary antibodies, signals were analyzed with HRP substrate and detected by Gel Imaging system (Tanon).

## In-vitro GST pull-down assay

0.1 mg GST-tagged bait protein was captured with GS4B resin for 30 min at 4 and supernatant was removed by 800 g centrifugation. A small portion of sample was set aside as input and analyzed by SDS-PAGE. Protein samples for analysis were mixed with rest pre-incubated beads for 2 h at 4 □. After three cycles wash with lysis buffer plus 0.02% (w/v) DDM, samples were extracted with 2x SDS-loading buffer and analyzed by immunoblotting.

## Co-immunoprecipitation assay

Cells were transfected as described above. After transfection, cells were harvested by centrifugation at 800 g and washed twice with PBS, followed by lysing with Co-IP buffer containing 50 mM Tris pH 7.4, 150 mM NaCl, 1 mM EDTA, protease inhibitor cocktail and detergent (0.5% (v/v) NP-40, 1% (w/v) DDM or 1% (w/v) GDN). NP-40 was selected if not mentioned. The supernatant after 20,000 g centrifugation was collected and incubated with HA-Nanoab-Agarose beads. The beads were further washed three times with Co-IP buffer. Proteins were extracted with 2x SDS-loading buffer and analyzed by immunoblotting.

## Generation of CRISPR KO cell lines using lentivirus

HEK293T cells were used to produce lentiviruses. In brief, 14 μg lentiviral plasmids, together with packaging plasmids including 6 μg psPAX2 and 4 μg pMD2.G were mixed with 150 μg polyethylenimine (PEI) for HEK293T transfection in a 10-cm dish. After 12 h transfection, medium was exchanged to RPMI 1640 with 10% (v/v) FBS, antibiotics (100 U/ml penicillin, 0.1 mg/ml streptomycin) and 0.05 mM β-ME. Then transfected HEK293T cells were continued to culture for 48 h. Then supernatants containing lentiviruses were collected by centrifugation, filtered via 0.22-μm polyethersulfone (PES) filters and added into THP1 cells. Cells were infected for 48 h in incubator, and then selected with the respective antibiotics.

## RNA sequencing

For sample preparation, 5×10^6^ THP1 cells per condition were stimulated with 20 nM pro-IL-1β, 20 nM IL-1β or left untreated (n=3). After 16 h treatment, cells were washed twice with RNase-free PBS and lysed with 1 mL Trizol. Total RNA was extracted using the standard protocol. 1 μg total RNA was used for the following library preparation. For mRNA enrichment, poly(A) mRNA isolation was performed using Oligo(dT) beads. Alternatively, when the sample quality or experimental design necessitated, ribosomal RNA was depleted using rRNA removal methods to enrich for coding and non-polyadenylated transcripts. The mRNA fragmentation was performed using divalent cations and high temperature. Priming was performed using Random Primers. First strand cDNA synthesis was carried out using reverse transcriptase, followed by second-strand cDNA synthesis. For strand-specific library construction, during the second-strand synthesis the standard dTTP was replaced with dUTP; subsequent treatment with USER enzyme selectively degrades the second strand prior to PCR amplification, thereby preserving strand information. The purified double-stranded cDNA was then treated to repair both ends and add a dA-tail in a single reaction, followed by a T-A ligation to add adaptors to both ends. Size selection of adaptor-ligated DNA was then performed using DNA Clean Beads. Each sample was then amplified by PCR using P5 and P7 primers, and the PCR products were validated. Finally, libraries with different indexes were multiplexed and loaded on an Illumina HiSeq instrument for sequencing using a 2×150 paired-end (PE) configuration according to the manufacturer’s instructions.

## Data analysis of RNA-seq

Raw sequencing reads in FASTQ format were processed using Fastp^30^ (v0.24.1) to remove technical sequences and low-quality bases, with the parameter of Phred quality cutoff = 20, error rate = 10%, Minimum read length = 75 bps and Maximum number of ambiguous (N) bases = 5. Read quality was confirmed using FastQC^31^ (v0.11.9) to ensure compliance with standard metrics. Clean reads were then aligned to the reference genome (*Homo sapiens*, GRCh38.p13) using HISAT2^32^ (v2.2.1), a splice-aware aligner. The reference genome and annotation files were obtained from ENSEMBL. Gene-level counts were quantified using HTSeq^33^ (v0.6.1). Reads overlapping exons were counted, and multi-mapping reads or reads overlapping multiple genes were excluded. DEGs were identified using DESeq2^34^ (v1.34.0), which employs a negative binomial distribution model. Genes with |log2FC| ≥ 1 and FDR < 0.05 were classified as differentially expressed, if not provided otherwise. Overrepresented terms were identified using GOSeq^35^ (v1.34.1) with the Wallenius approximation to correct for gene length bias, and GO terms were visualized using topGO.

## Multiple sequence alignments

Full-length human TLR paralog sequences were aligned with Clustal Omega Program^36^ on Uniprot (https://www.uniprot.org/align). The following sequences were used: TLR1 (Uniprot: Q15399), TLR2 (Uniprot: O60603), TLR3 (Uniprot: O15455), TLR4 (Uniprot: O00206), TLR5 (Uniprot: O60602), TLR6 (Uniprot: Q9Y2C9), TLR7 (Uniprot: Q9NYK1), TLR8 (Uniprot: Q9NR97), TLR9 (Uniprot: Q9NR96) and TLR10 (Uniprot: Q9BXR5).

## Data processing and statistics

All statistics in the current study represented Mean± SEM, two-way ANOVA and unpaired student t-test performed, ns = p>0.05, * = p ≤0.05, ** = p ≤0.01, *** = p ≤0.001, **** = p ≤ 0.0001. Data processing and figures preparation used Graphpad prism 10 (GraphPad Software, Boston, Massachusetts USA, www.graphpad.com).

## Legends for Supplementary Figures

**Supplementary Figure 1.**
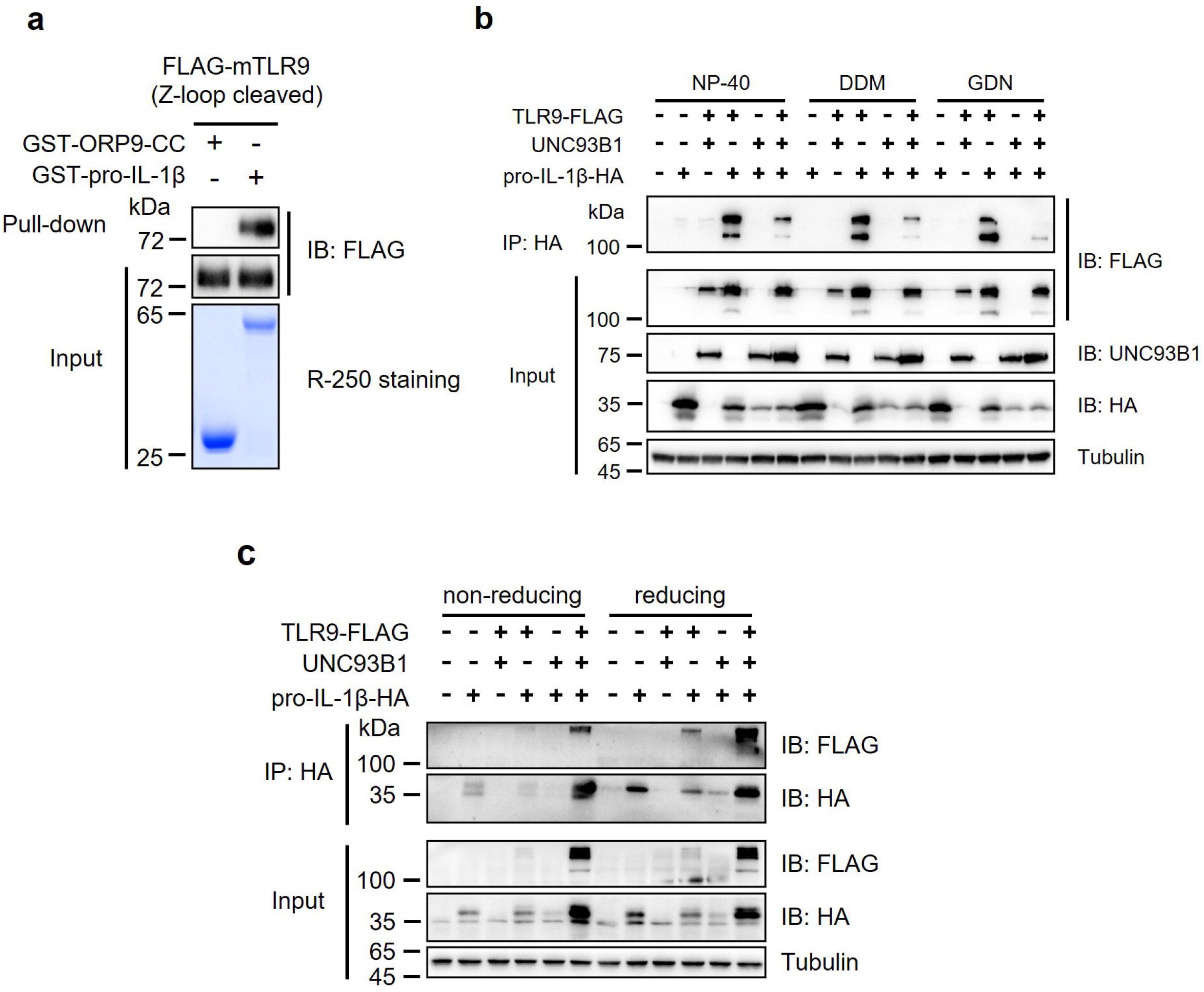
| Discovery of the interaction between pro-IL-1β and TLR9. **a,** GST pull-down of FLAG-mTLR9 (mouse) and GST-tagged effectors. **b,** Immunoprecipitation assay of pro-IL-1β and TLR9 in different detergents. HEK293T cells were transfected with TLR9, UNC93B1, pro-IL-1β or plasmid combinations. After 48 h transfection, cells were lysed with Co-IP buffer containing different detergents and analyzed with immunoblotting. **c,** Immunoprecipitation assay and TLR9 in non-reducing and reducing conditions. Protocol of co-IP is the same as NP-40 group in panel b, except that 5 mM β-ME was added throughout the process in reducing groups.

**Supplementary Figure 2.**
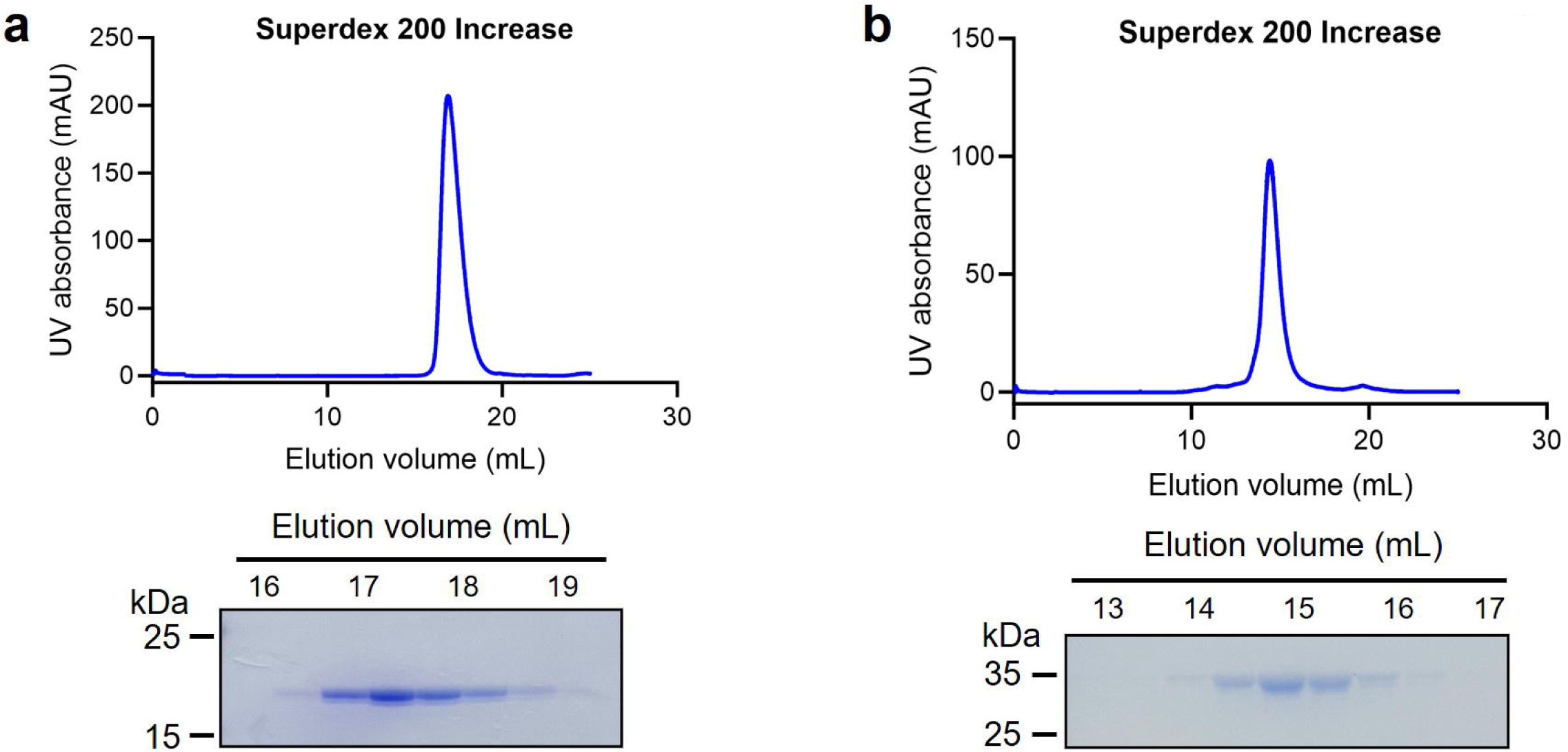
| Purification of recombinant human pro-IL-1β and IL-1β. **a,** Size-exclusion chromatography of human IL-1β. **b,** Size exclusion chromatography of human pro-IL-1β.

**Supplementary Figure 3.**
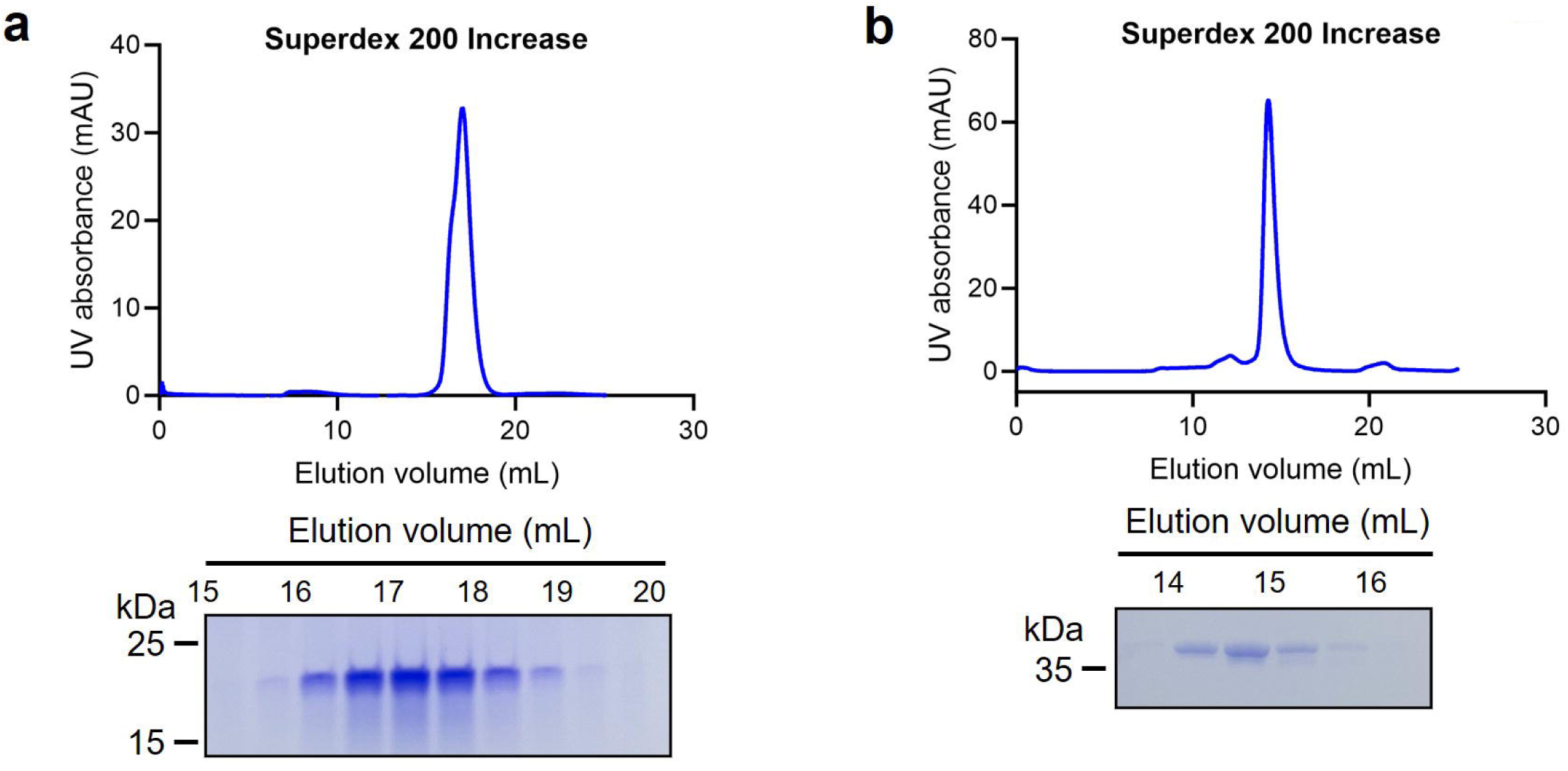
| Purification of recombinant mouse pro-IL-1β and IL-1β. **a,** Size-exclusion chromatography of mouse IL-1β. **b,** Size exclusion chromatography of mouse pro-IL-1β mutant (C9S, C34S, C42S, C104S, C116S).

**Supplementary Figure 4.**
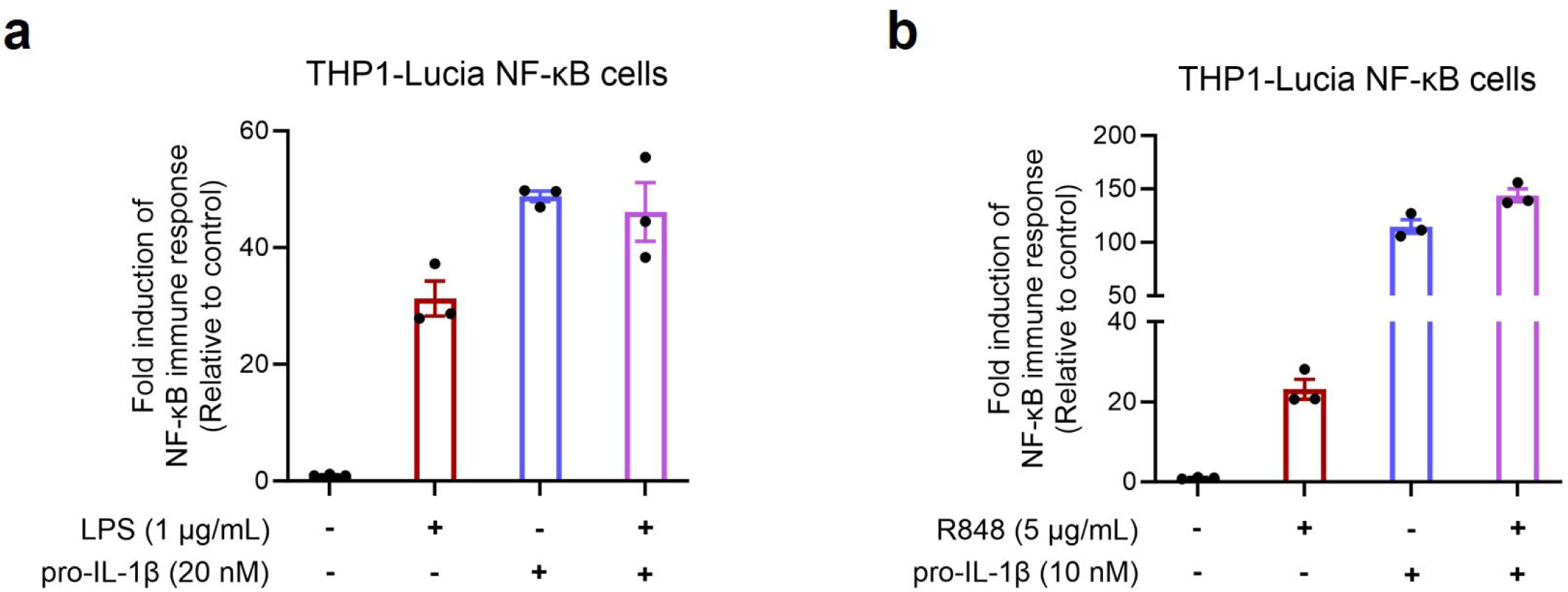
| Synergy analysis of pro-IL-1β with LPS or R848. THP1-Lucia NF-κB cells was treated with pro-IL-1β/LPS (**a**) or pro-IL-1β/R848 (**b**). Medium were harvested after 16 h for luciferase assay.

**Supplementary Figure 5.**
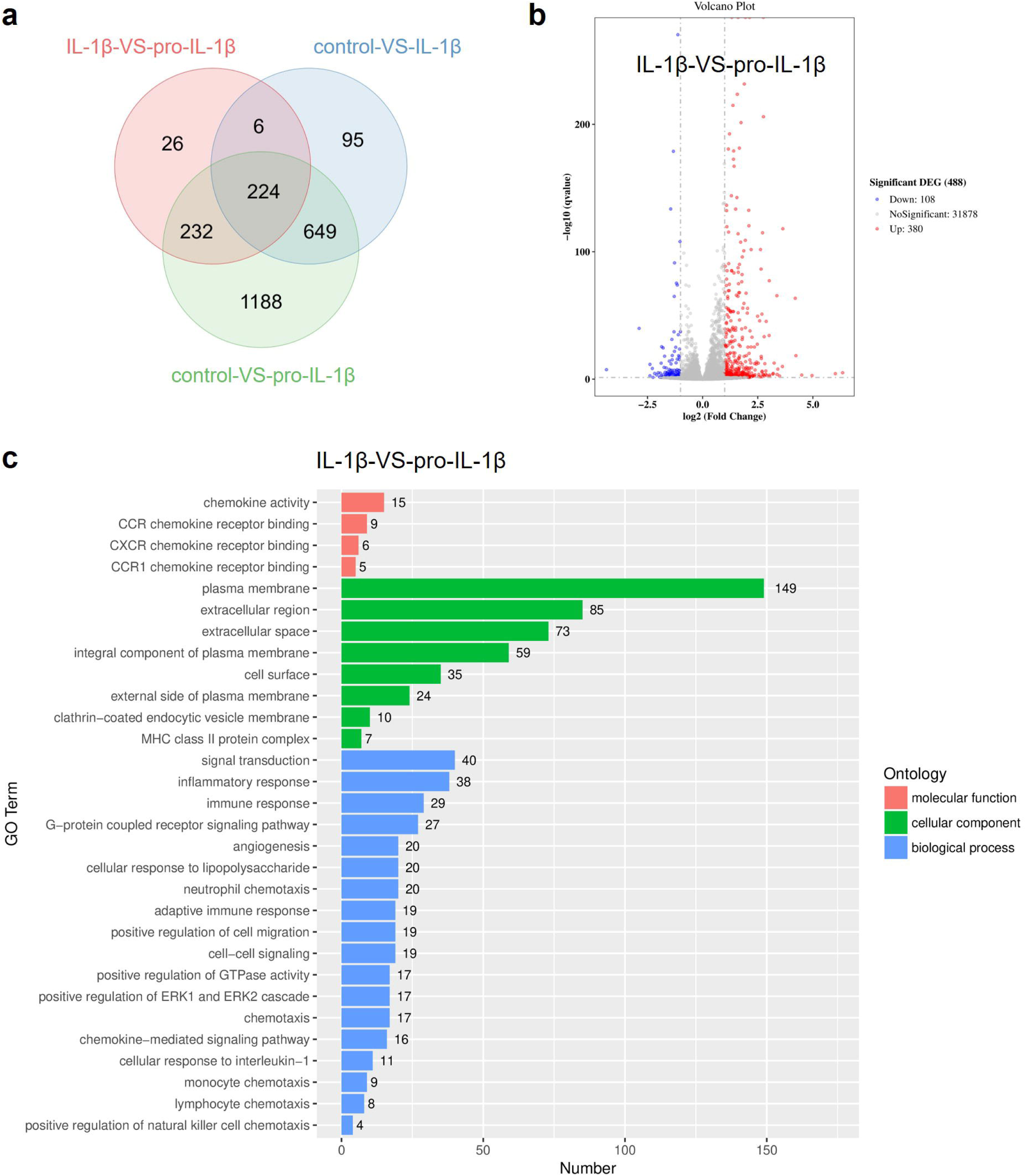
| RNA-seq analysis of untreated and cytokine-treated groups. THP1 cells were treated with 20 nM IL-1β or 20 nM pro-IL-1β (n = 3, 5×10^6^ cells for each group). Cells were stimulated for 16 h, washed, lysed with Trizol and delivered for RNA-seq. **a,** Venn diagram of control, IL-1β treatment and pro-IL-1β treatment. Blue circle represents genes regulated by IL-1β alone, as green circle represents genes regulated by pro-IL-1β alone. Light pink circle represents genes regulated by pro-IL-1β compared with IL-1β. **b,** Volcano plot for presentation of the gene regulation differences between IL-1β and pro-IL-1β. Threshold of differential genes: |Log_2_ (Fold change)|≥1 and q_value_≤0.05. **c,** Biological process that pro-IL-1β is involved in, presented as GO term diagram. X-axis stands for the numbers of differential genes in each term, and y-axis stands for the GO term.

**Supplementary Figure 6.**
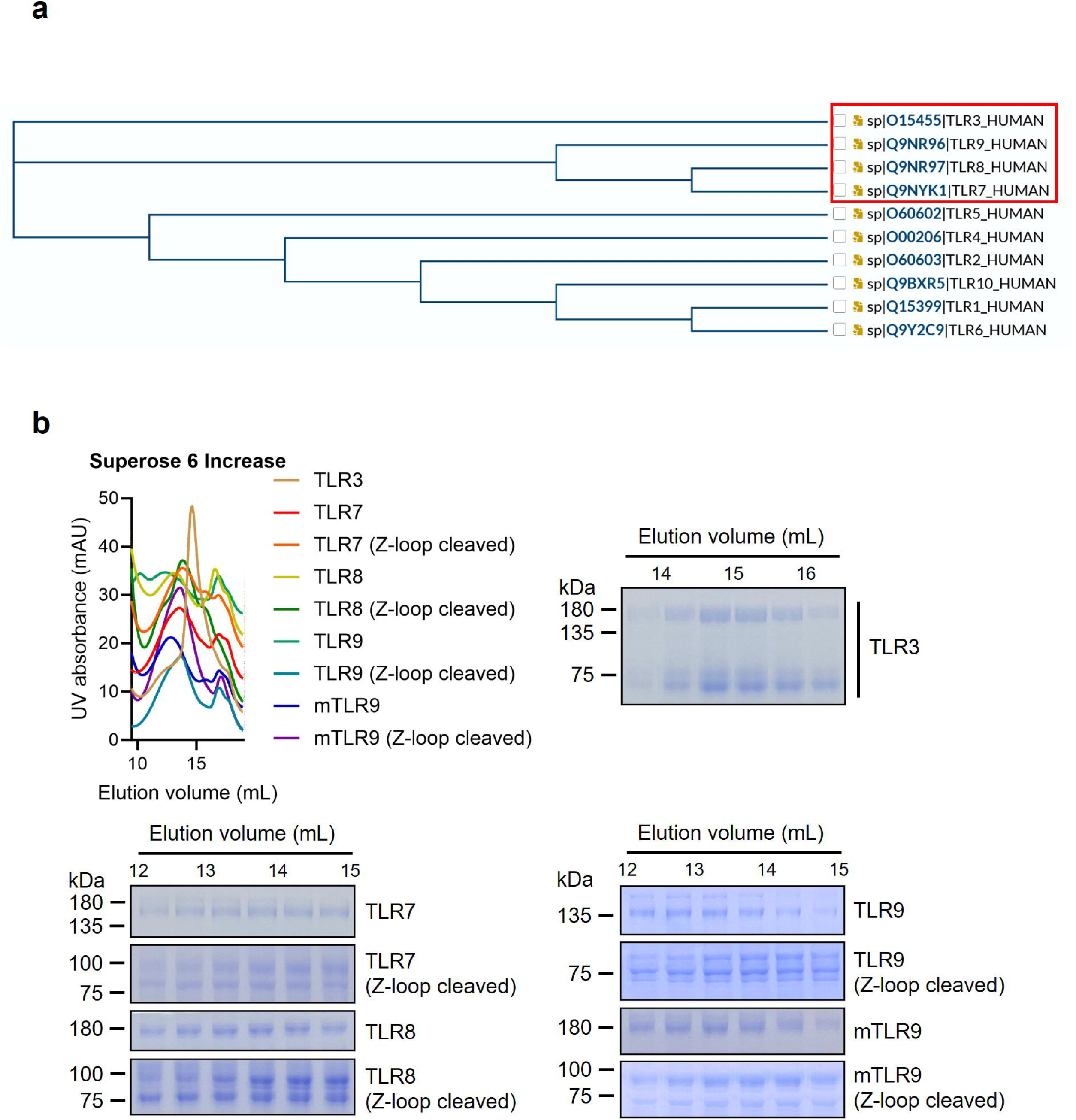
| Purification of Toll-like receptors. **a**, Phylogenetic tree of human TLRs, aligned using Uniprot (https://www.uniprot.org). **b,** Size-exclusion chromatography of TLRs. mTLR9: mouse TLR9.

**Supplementary Figure 7.**
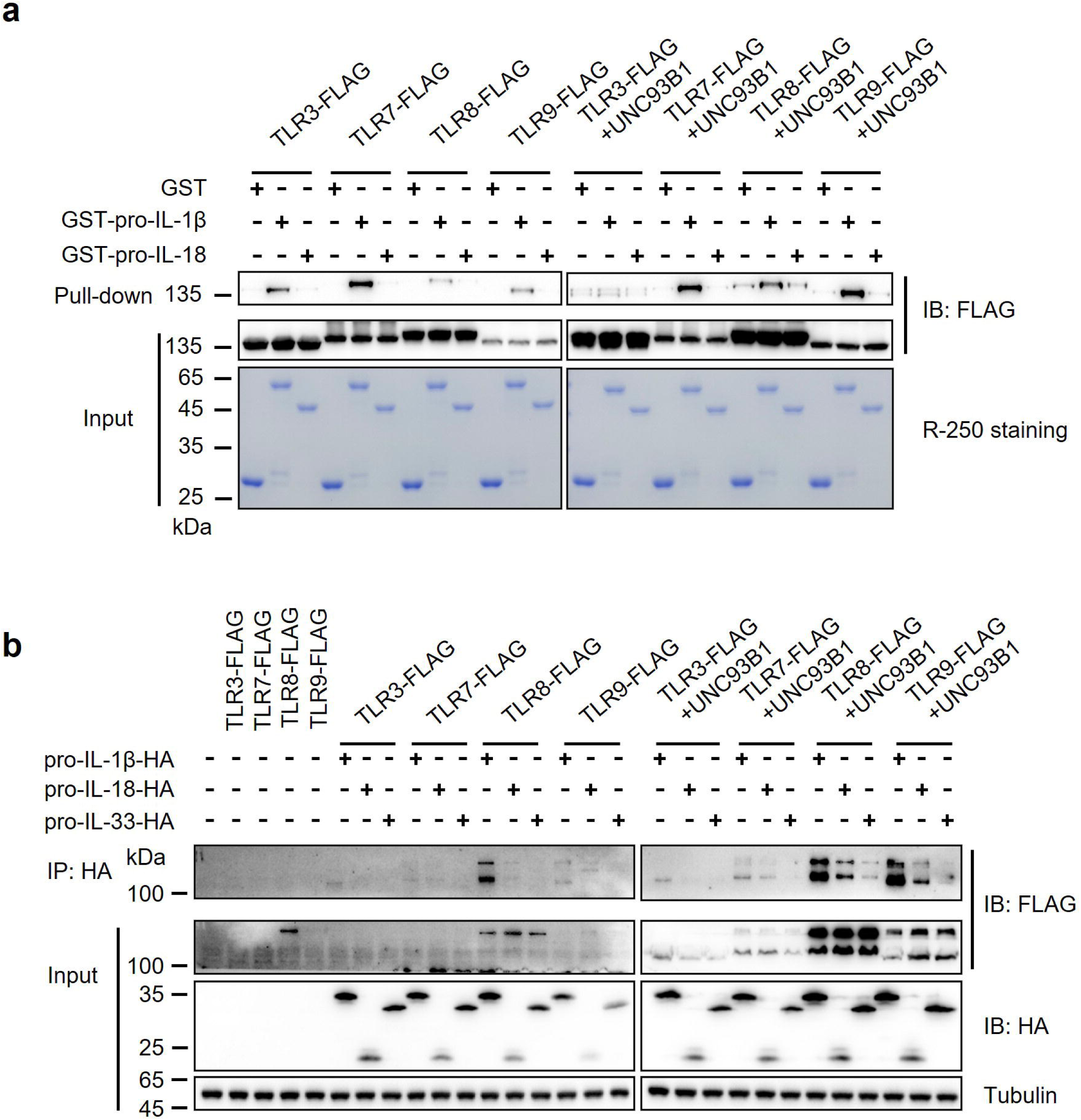
| pro-IL-1β binds to TLR3/7/8/9 in vitro. **a,** In vitro pull-down of purified TLR3/7/8/9 and cytokines. **b,** Co-IP of TLR3/7/8/9 and cytokines. HEK293T cells were used for overexpression. After 48 h transfection, cells were lysed and the supernatants were applied for immunoprecipitation.

