## Supplementary Table 1 for "Pro-IL-1β activates NF-κB independently of maturation"

**Table S1 Endotoxin concentration of target proteins**

| Protein name | Conversion of protein concentration | Endotoxin concentration *（EU/μg） |
| --- | --- | --- |
| IL-1β | 1 μM = 17 μg/mL | 0.067 |
| pro-IL-1β | 1 μM = 31 μg/mL | 0.060 |
| mIL-1β | 1 μM = 17 μg/mL | 0.023 |
| mpro-IL-1β (mut) | 1 μM = 31 μg/mL | 0.56 |
| IL-1Ra | 1 μM = 17 μg/mL | 0.018 |
| pro-peptide | 1 μM = 14 μg/mL | 0.90 |

* The concentration value of endotoxin is retained to two significant figures.
